# RefPlantNLR: a comprehensive collection of experimentally validated plant NLRs

**DOI:** 10.1101/2020.07.08.193961

**Authors:** Jiorgos Kourelis, Toshiyuki Sakai, Hiroaki Adachi, Sophien Kamoun

## Abstract

Reference datasets are critical in computational biology. They help define canonical biological features and are essential for benchmarking studies. Here, we describe a comprehensive reference dataset of experimentally validated plant NLR immune receptors. RefPlantNLR consists of 442 NLRs from 31 genera belonging to 11 orders of flowering plants. This reference dataset has several applications. We used RefPlantNLR to determine the canonical features of functionally validated plant NLRs and to benchmark the five most popular NLR annotation tools. This revealed that although NLR annotation tools tend to retrieve the majority of NLRs, they frequently produce domain architectures that are inconsistent with the RefPlantNLR annotation. Guided by this analysis, we developed a new pipeline, NLRtracker, which extracts and annotates NLRs based on the core features found in the RefPlantNLR dataset. The RefPlantNLR dataset should also prove useful for guiding comparative analyses of NLRs across the wide spectrum of plant diversity and identifying under-studied taxa. We hope that the RefPlantNLR resource will contribute to moving the field beyond a uniform view of NLR structure and function.

## INTRODUCTION

Reference datasets are critical in computational biology (Weber *et al*., 2019; Schaafsma and Vihinen, 2018). They help define canonical biological features and are essential to benchmarking studies. Reference datasets are particularly important for defining the sequence and domain features of gene and protein families. Despite this, curated collections of experimentally validated sequences are still lacking for several widely studied gene and protein families. One example is the nucleotide-binding leucine-rich repeat (NLR) family of plant proteins. NLRs constitute the predominant class of disease resistance (*R*) genes in plants (Jones *et al*., 2016; Kourelis and van der Hoorn, 2018; de Araújo *et al*., 2020). They function as intracellular receptors that detect pathogens and activate an immune response that generally leads to disease resistance. NLRs are thought to be engaged in a coevolutionary tug of war with pathogens and pests. As such, they tend to be among the most polymorphic genes in plant genomes, both in terms of sequence diversity and copy-number variation (Tamborski and Krasileva, 2020). Ever since their first discovery in the 1990s, hundreds of NLRs have been characterized and implicated in pathogen and self-induced immune responses (Kourelis and van der Hoorn, 2018). NLRs are among the most widely studied and economically valuable plant proteins given their importance in breeding crops with disease resistance (Dangl *et al*., 2013).

NLRs occur widely across all kingdoms of life where they generally function in non-self perception and innate immunity (Jones *et al*., 2016; Uehling *et al*., 2017; Gao *et al*., 2020). In the broadest biochemical definition, NLRs share a similar multidomain architecture consisting of a nucleotide-binding and oligomerization domain (NOD) and a super-structure forming repeat (SSFR) domain (Dyrka *et al*., 2020). The NOD is either an NB-ARC (nucleotide-binding adaptor shared by APAF-1, certain *R* gene products and CED-4) or NACHT (neuronal apoptosis inhibitory protein, MHC class II transcription activator, HET-E incompatibility locus protein from *Podospora anserina*, and telomerase-associated protein 1), whereas the SSFR domain can be formed by ankyrin (ANK) repeats, tetratricopeptide repeats (TPRs), armadillo (ARM) repeats, WD repeats or leucine-rich repeats (LRRs) (Dyrka *et al*., 2020; Andolfo *et al*., 2019). Plant NLRs exclusively carry an NB-ARC domain with the C-terminal SSFR consisting typically of LRRs. The NB-ARC domain has been used to determine the evolutionary relationships between plant NLRs given that it is the only domain that produces reasonably good global alignments across all members of the family. In flowering plants (angiosperms), NLRs form three main monophyletic groups with distinct N-terminal domain fusions: the TIR-NLR subclade containing an N-terminal Toll/interleukin-1 receptor (TIR) domain, the CC-NLR-subclade containing an N-terminal Rx-type coiled-coil (CC) domain and the CC_R_-NLR subclade containing an N-terminal RPW8-type CC (CC_R_) domain (Shao *et al*., 2016). Additionally, Lee *et al*. (2020) have recently proposed that the G10-subclade of NLRs is a monophyletic group containing a distinct type of CC (here referred to as CC_G10_; CC_G10_-NLR). NLRs also occur in non-flowering plants where they carry additional types of N-terminal domains such as kinases and α/β hydrolases (Andolfo *et al*., 2019).

Plant NLRs likely evolved from multifunctional receptors to specialized receptor pairs and networks (Adachi, Contreras, *et al*., 2019; Adachi, Derevnina, *et al*., 2019). NLRs which combine pathogen detection and immune signalling activities into a single protein are referred to as “functional singletons”, whereas NLRs which have specialized in pathogen recognition or immune signalling are referred to as “sensor” or “helper” NLRs, respectively. About one quarter of NLR genes occur as “genetic singletons” in plant genomes, whereas the others form genetic clusters often near telomeres (Jacob *et al*., 2013). This genomic clustering likely aids the evolutionary diversification of this gene family and subsequent emergence of pairs and networks (Tamborski and Krasileva, 2020; Adachi, Derevnina, *et al*., 2019). The emerging picture is that NLRs form genetic and functional receptor networks of varying complexity (Wu *et al*., 2018; Adachi, Derevnina, *et al*., 2019).

The mechanism of pathogen detection by NLRs can be either direct or indirect (Kourelis and van der Hoorn, 2018). Direct recognition involves the NLR protein binding a pathogen-derived molecule or serving as a substrate for the enzymatic activity of a pathogen virulence protein (known as effectors). Indirect detection is conceptualized by the guard and decoy models where the status of a host component–the guardee or decoy–is monitored by the NLR (van der Biezen and Jones, 1998; van der Hoorn and Kamoun, 2008). Some sensor NLRs known as NLR-IDs contain non-canonical “integrated domains” that can function as decoys to bait pathogen effectors and enable pathogen detection (Cesari *et al*., 2014; Sarris *et al*., 2016; Wu *et al*., 2015). These extraneous domains appear to have evolved by fusion of an effector target domain into an NLR (Cesari *et al*., 2014; Sarris *et al*., 2016; Białas *et al*., 2017). The sequence diversity of integrated domains in NLR-IDs is staggering indicating that novel domain acquisitions have repeatedly occurred throughout the evolution of plant NLRs (Sarris *et al*., 2016; Kroj *et al*., 2016).

Given their multidomain nature, sequence diversity and complex evolutionary history, prediction of NLR genes from plant genomes is challenging. Several bioinformatic tools have been developed to extract plant NLRs from sequence datasets. As an input these tools take either annotated genomic features and transcriptomic data, or alternatively can be run directly on the unannotated genomic sequence. NLR-Parser, RGAugury, RRGPredictor, and DRAGO2 identify transcript and protein sequences that have features of NLRs and are best described as NLR extractors (Steuernagel *et al*., 2015; Li *et al*., 2016; Osuna-Cruz *et al*., 2018; Santana Silva and Micheli, 2020). RGAugury, RRGPredictor, and DRAGO2 also extract other classes of immune-related genes in addition to NLRs. These various tools use pre-defined motifs to classify sequences as NLRs, but they differ in the methods and pipelines. NLR-Annotator –an extension of NLR-Parser–and NLGenomeSweeper, can also use unannotated genome sequences as input to predict the genomic locations of NLRs (Steuernagel *et al*., 2020; Toda *et al*., 2020). This output then requires manual annotation to extract the final gene-models and some of the annotated loci may represent partial or pseudogenized genes.

The goal of this study is to provide a curated reference dataset of experimentally validated plant NLRs. This version of RefPlantNLR (v.20201113_442) consists of 442 NLRs from 31 genera belonging to 11 orders of flowering plants. We used RefPlantNLR to determine the canonical features of functionally validated plant NLRs and benchmark NLR annotation tools. We found that these NLR-annotation tools are able to extract the majority of NLRs in the RefPlantNLR dataset, however, the domain architecture analysis produced by these tools is often inconsistent with that of RefPlantNLR. In order to simplify NLR extraction, annotation, and phylogenetic analysis we developed NLRtracker: a pipeline that uses InterProScan (Finn *et al*., 2017) and predefined NLR motifs (Jupe *et al*., 2012) to extract NLRs and provide domain architecture analyses based on the canonical features found in the RefPlantNLR dataset. Additionally, NLRtracker outputs the extracted NB-ARC domain facilitating downstream phylogenetic analysis. RefPlantNLR should also prove useful in guiding comparative and phylogenetic analyses of plant NLRs and identifying under-studied taxa for future studies.

## RESULTS and DISCUSSION

### RefPlantNLR pipeline

The current version of RefPlantNLR (v.20201113_442) contains 442 entries from 31 genera of flowering plants (**Figure 1**; **Supplemental dataset 1**, **2**, **3**). The list was obtained by mining and assessing information gathered from the literature. Briefly, we manually crawled through the literature, extracting plant NLRs that have been experimentally validated to at least some degree. We defined experimental validation broadly as genes reported to be involved in any of the following: 1) disease resistance, 2) disease susceptibility, including effector-triggered immune pathology or trailing necrosis to viruses, 3) hybrid necrosis, 4) autoimmunity, 5) NLR helper function or involvement in downstream immune responses, 6) negative regulation of immunity, and 7) well-described allelic series of NLRs with different pathogen recognition spectra even if not reported to confer disease resistance.

**Figure 1:**
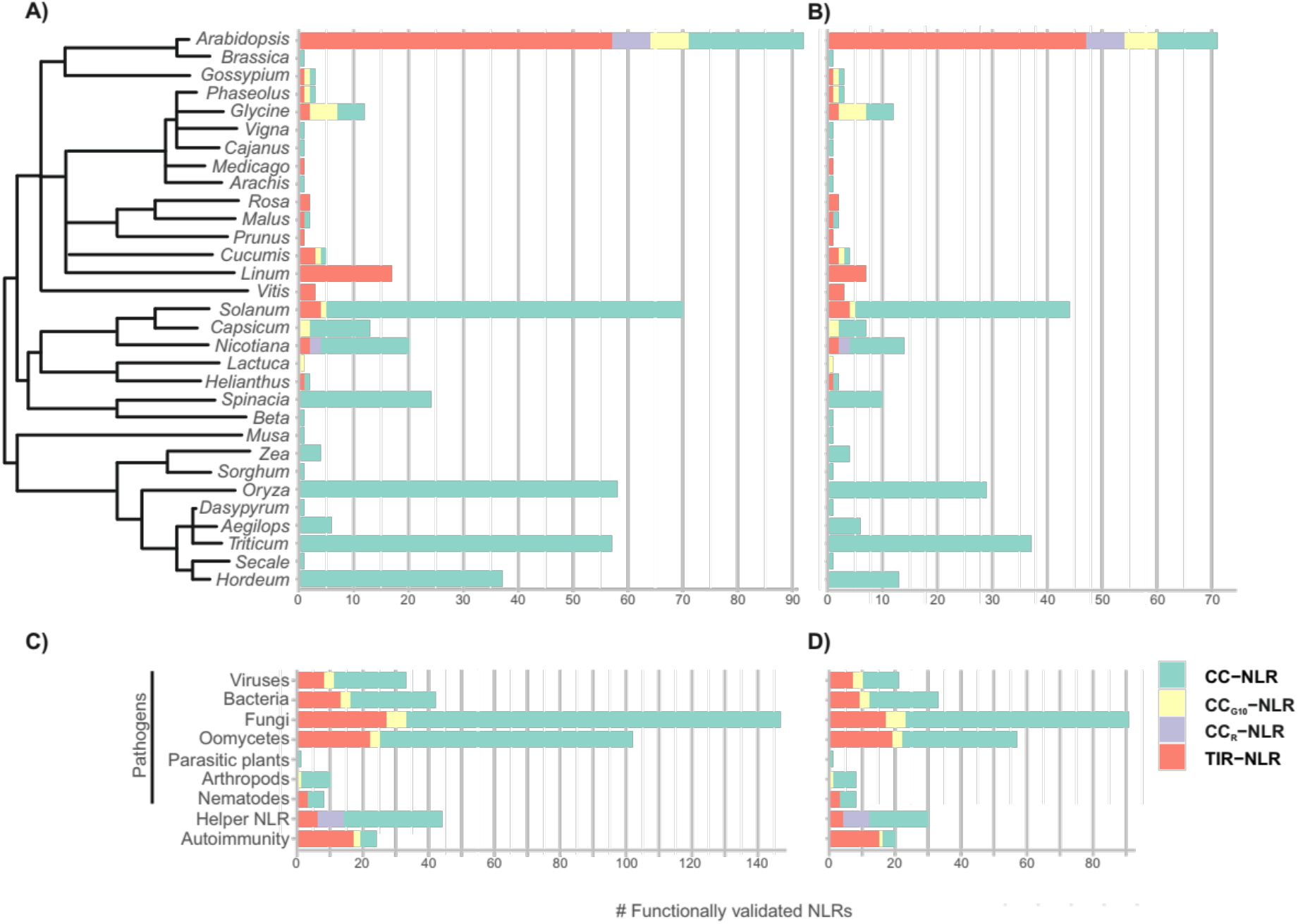
Number of experimentally validated RefPlantNLR sequences per plant genus. **A)** The number of experimentally validated NLRs per plant genus (N = 442), and **B)** the per genus reduced redundancy set at a 90% sequence similarity threshold (N = 285) are plotted as a stacked bar graph. **C)** The class of pathogen to which NLRs in the RefPlantNLR dataset confer a response. Some NLRs may be involved in the response against multiple classes of pathogens while others have a helper role or are found to be involved in allelic variation in autoimmune/hybrid necrosis responses, and **D)** the per genus reduced redundancy set at a 90% sequence similarity threshold are plotted as a stacked bar graph. The number of experimentally validated NLRs belonging to the monophyletic TIR-NLR, CC-NLR, CC_R_-NLR, or CC_G10_-NLR subclade members are indicated.

To validate the recovered sequences as genuine NLRs, we annotated them using InterProScan (Finn *et al*., 2017; see Material & Methods for the used sequence signatures) and predefined NLR motifs (Jupe *et al*., 2012). We defined NLRs as sequences containing the NB-ARC domain (Pfam signature PF00931) or a P-loop containing nucleoside triphosphate hydrolases domain (SUPERFAMILY signature SSF52540) combined with plant-specific NLR motifs (see Material & Methods for the used motifs). This resulted in 441 sequences. We also included RXL, which carries C-terminal LRRs, as well as an N-terminal Rx-type CC EDVID motif and the CC-type RNBS-D motif of the NB-ARC domain, but otherwise does not get annotated with a P-loop containing nucleoside triphosphate hydrolase domain (Xie *et al*., 2020). Altogether these 442 sequences form the current version of RefPlantNLR (**Table S1**).

In addition to the 442 NLRs present in this version of RefPlantNLR, we separately collected several characterized animal, bacterial, and archaeal NB-ARC proteins (**Table S2**, **Supplemental dataset 4**) which can be used as outgroups for comparative analyses. Furthermore, several characterized plant immune components have features often found in NLRs–such as the RPW8-type CC or the TIR domain–but lack the NB-ARC domain or NB-ARC-associated motifs that we used to define NLRs (see above). Since these proteins may have common origins with plant NLRs or may be useful for comparative analysis of these domains we have collected them separately as well (**Table S3**, **Supplemental dataset 5**, **6**, **7**).

### Description of the RefPlantNLR dataset

**Table S1** describes the RefPlantNLR dataset, including amino acid, coding sequence (CDS) and locus identifiers, as well as the organism from which the NLR was cloned, the article describing the identification of the NLR, the pathogen type and pathogen to which the NLR provides resistance (when applicable), the matching pathogen effector, additional host components required for pathogen recognition (guardees or decoys) or required for NLR function and the articles describing the identification of the pathogen and host components. We also provide domain annotations for the amino acid sequences (**Supplemental dataset 8**). From this dataset, we extracted 433 unique NLRs and 450 NB-ARC domains of which 370 were unique (**Supplemental dataset 9, 10**). NLRs with identical amino acid sequences were recovered because they have different resistance spectra when genetically linked to different sensor NLR allele (e.g. alleles of Pik), are different in non-coding regions leading to altered regulation (e.g. RPP7 alleles) or have been independently discovered in different plant genotypes (e.g. RRS1-R and SLH1).

The distribution of the RefPlantNLR entries across plant species mirrors the most heavily studied taxa, i.e. *Arabidopsis*, Solanaceae (*Solanum, Capsicum* and *Nicotiana*) and cereals (*Oryza, Triticum* and *Hordeum*) (**Figure 1A**). These seven genera comprise 79% (347 out of 442) of the RefPlantNLR sequences. When accounting for redundancy by collapsing similar sequences (>90% overall amino acid identity per genus), these seven genera would still account for 75% (215 out of 285) sequences (**Figure 1B**). It should be noted that there could be different evolutionary rates between NLRs, and hence some subfamilies may still be overrepresented in the reduced redundancy set.

In total, 31 plant genera representing 11 taxonomic orders are listed in RefPlantNLR. Interestingly, these species represent a small fraction of plant diversity with only 11 of 59 major seed plant (spermatophyte) orders described by Smith and Brown represented, and not a single entry from non-flowering plants (**Table S4**) (Smith and Brown, 2018). Arabidopsis remains the only species with experimentally validated NLRs from the four major clades (CC-NLR, CC_G10_-NLR, CC_R_-NLR and TIR-NLR) (**Figure 1**).

We also mapped the frequency of the pathogens that are targeted by RefPlantNLR entries. Most validated NLRs in the RefPlantNLR dataset are involved in responses against fungi followed by oomycetes (**Figure 1c**, **Figure 1d** for the reduced redundancy set). Responses to certain pathogen taxa is not constrained to particular subclasses of NLRs as all of TIR-NLRs, CC_G10_-NLRs, and CC-NLRs are involved in resistance to the main pathogen classes (fungi, oomycete, bacteria, and viruses). The notable exception is the CC_R_-NLR subclade which has only been validated for its helper function (**Figure 1c**, **Figure 1d**). Additionally, CC_G10_-NLR subclade members have not been assigned a helper activity, and CC_R_-NLR subclade members have not been implicated in autoimmunity or hybrid necrosis (**Figure 1c**, **Figure 1d**) even though several RPW8-only proteins are involved in hybrid necrosis (Chae *et al*., 2014; Barragan *et al*., 2019).

The average length of RefPlantNLR sequences varies depending on the subclass (**Figure 2A**, **Figure 2C** for the reduced redundancy set). CC-NLRs varied from 665 to 1845 amino acids (mean = 1021, N = 318), whereas TIR-NLR varied from 380 to 2048 amino acids (mean = 1167, N = 96). NB-ARC domains were more constrained (mean = 271, N = 370, stdev = 17) (**Figure 2B**). Nonetheless, 13 atypically short NB-ARCs (166 to 231 amino acids) and 5 long NB-ARCs (305 to 339 amino acids) were observed at more than two standard deviations of the mean illustrating the overall flexibility of plant NLRs even for this canonical domain (**Figure 2B, Figure 2D** for the reduced redundancy set).

**Figure 2:**
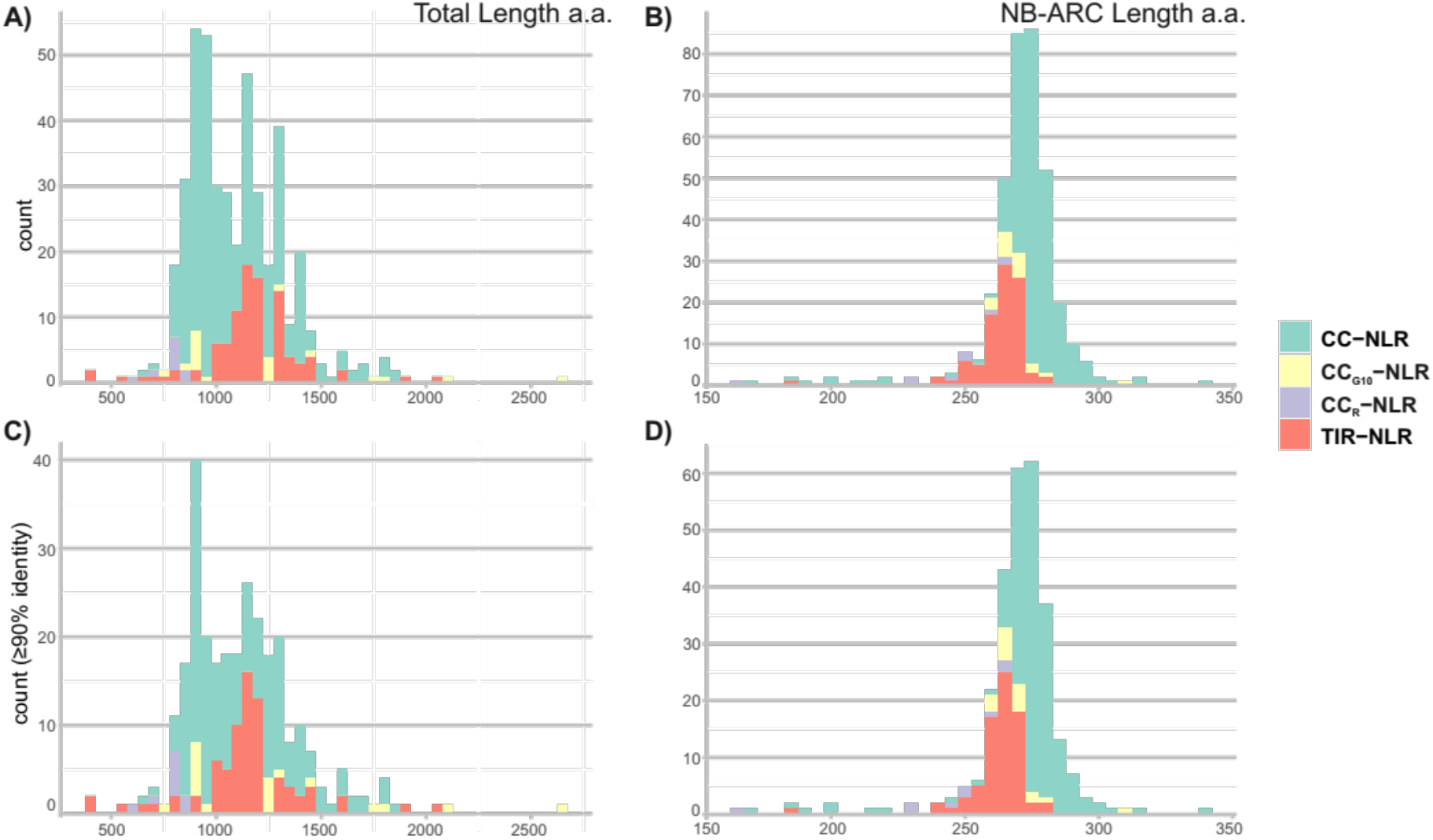
Length distribution RefPlantNLR amino acid sequence and extracted NB-ARC domains. Length distribution of the RefPlantNLR sequences. **A)** Histogram of RefPlantNLR amino acid sequence length (binwidth 50aa, N = 442). **B)** Histogram of the unique RefPlantNLR extracted NB-ARC domain (SUPERFAMILY signature SSF52540) amino acid sequence length (binwidth 5aa, N = 370). **C)** Histogram of amino acid sequence length of the reduced redundancy RefPlantNLR set at a 90% amino acid similarity threshold (binwidth 50aa, N = 285). **D)** Histogram of the extracted NB-ARC domain from the reduced redundancy RefPlantNLR set (binwidth 5aa, N = 281). Colour-coding according to NLR subfamily.

We noted that some of the unusually small NLRs lacked a SSFR domain, while some of the small NB-ARC domains appeared to be partial duplications of this domain. In order to look at domain architecture of NLRs more widely and to determine whether these unusual features are common, we mapped the domain architecture of RefPlantNLR proteins (**Figure 3A, Figure 3B** for the reduced redundancy set). Even though CC-NLR and TIR-NLR domain combinations were the most frequent (60% and 18%, respectively), we observed additional domain combinations. In the RefPlantNLR dataset, a subset of NLRs lack the N-terminal domain but still group with the major NLR clades based on the NB-ARC phylogeny. Some TIR-NLRs lack a SSFR domain. Non-canonical integrated domains are found in all NLR subfamilies, and occur at the N-terminus, in between the N-terminal domain and the NB-ARC domain, at the C-terminus or both ends. Of these non-canonical domains, the N-terminal late-blight resistance protein R1 domain (also known as the Solanaceae domain; Pfam signature PF12061) only occurs in association with the NB-ARC domain, and has an ancient origin likely in the most recent common ancestor of the Asterids and Amaranthaceae (Seong *et al*., 2020). Other non-canonical domains are also more wide-spread, including the monocot-specific integration of a zinc-finger BED domain in between the CC and NB-ARC domain (Bailey *et al*., 2018; Marchal *et al*., 2018). Finally, some CC-NLRs have significantly truncated NB-ARC domain as is the case for Pb1 and RXL, the latter which using InterProScan only gets annotated with a LRR domain although it retains other NLR motifs such as the EDVID and RNBS-D motifs (**Figure 3C**).

**Figure 3:**
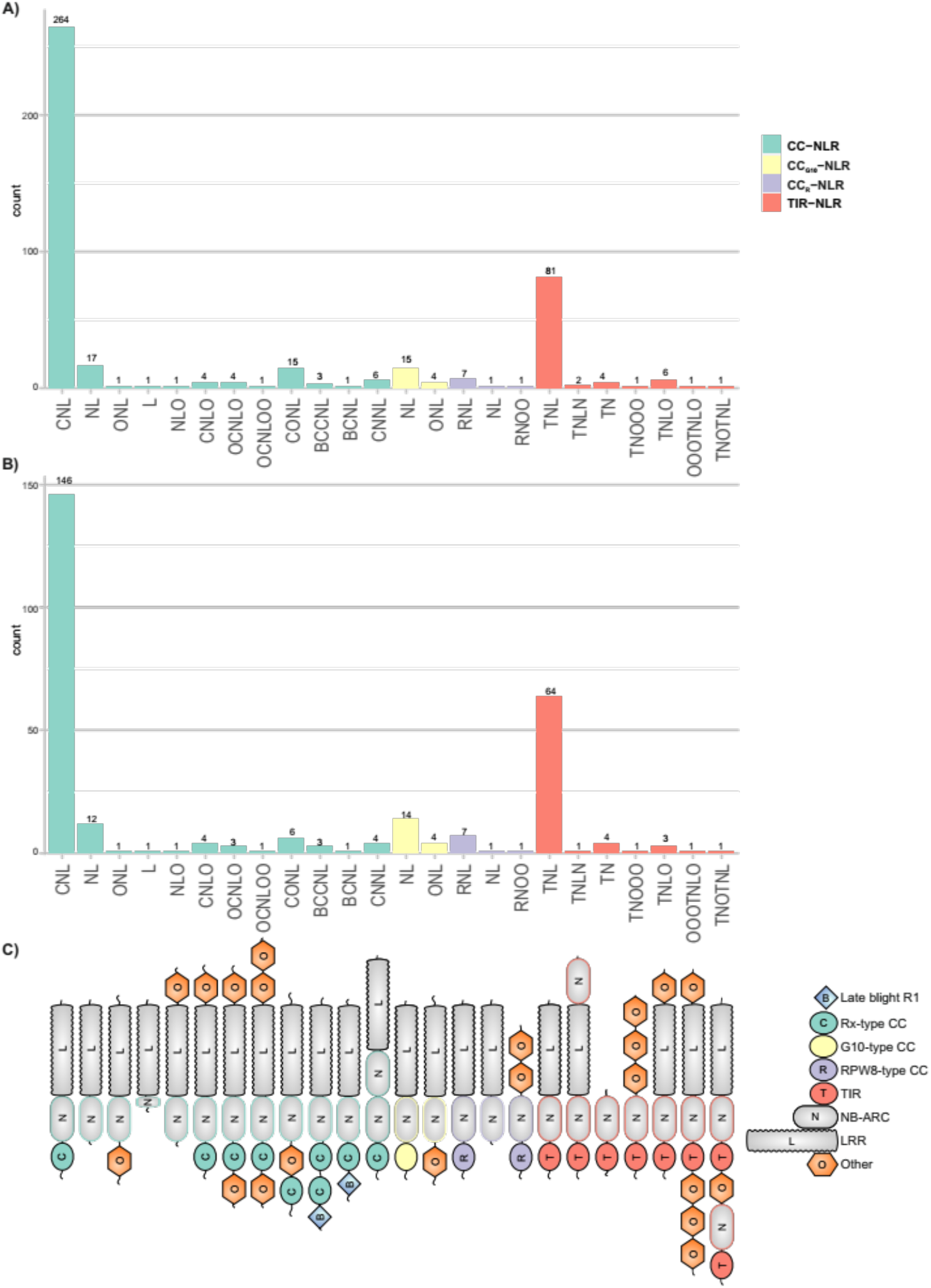
Domain architecture of the RefPlantNLRs. Bar chart of the domain architecture of **A)** RefPlantNLRs (N = 442), or **B)** the per genus reduced redundancy RefPlantNLR set at an overall 90% amino acid similarity per genus (N = 285). **C)** Schematic representation of domain architecture. Used InterPro signatures for each of the domains are highlighted in the material & methods. There is currently no InterProScan signature or motif for the CC_G10_ N-terminal domain.

We explored the phylogenetic diversity of RefPlantNLR proteins using the extracted NB-ARC domains (**Figure 4**; **Supplemental dataset 9, 10, 11, 12)**. As with previously reported NLR phylogenetic analyses, RefPlantNLR sequences generally grouped in well-defined clades, notably CC-NLR, CC_G10_-NLR, CC_R_-NLR and TIR-NLR. Within this phylogeny some of the branches, notably of Wed and Pi54, are long and may represent highly diverged NB-ARC domains. Since Pb1 (Hayashi *et al*., 2010) and RXL (Xie *et al*., 2020) do not match the Pfam NB-ARC domain they were not included in this phylogenetic analysis.

**Figure 4:**
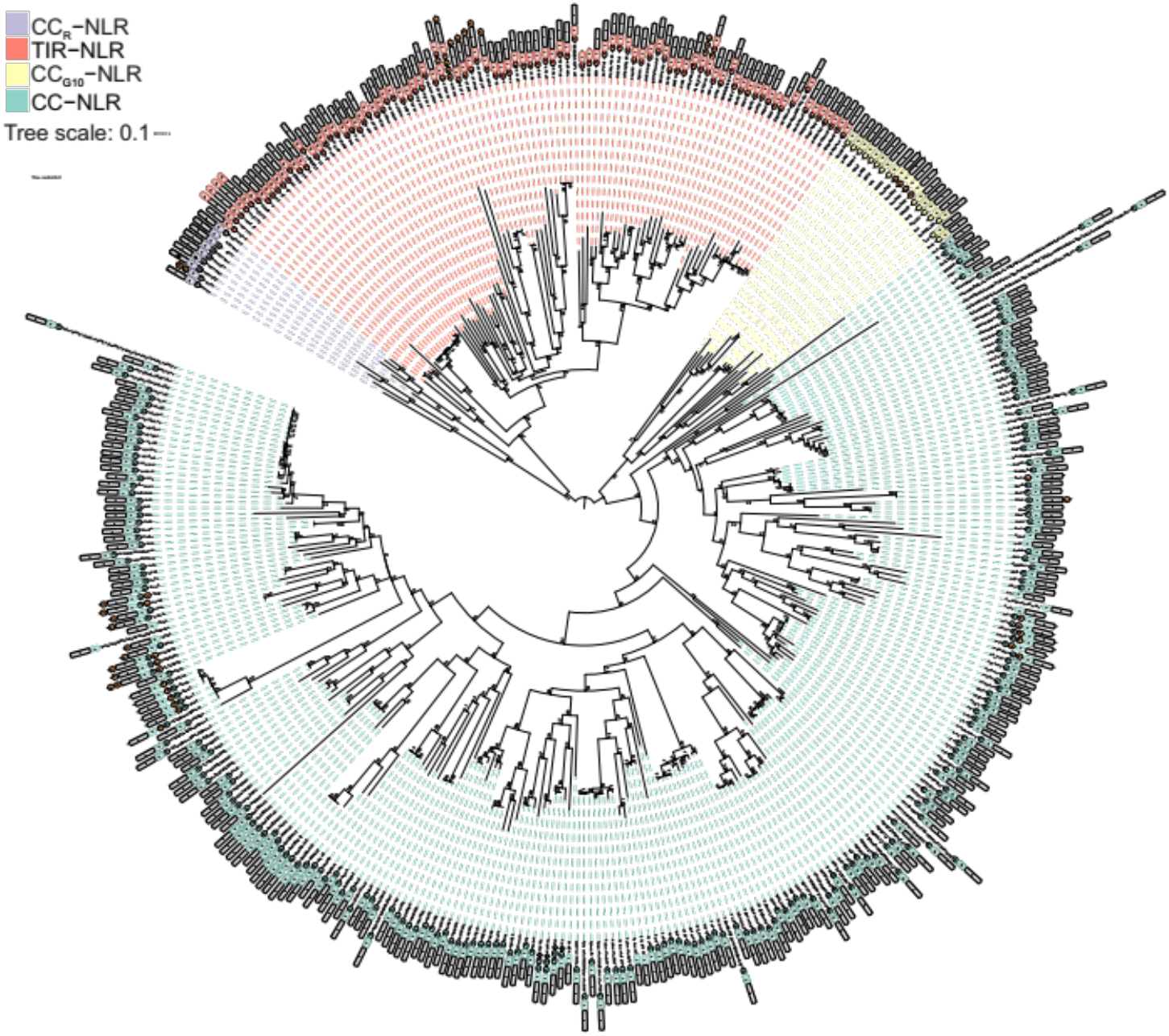
Phylogenetic diversity of RefPlantNLR sequences. The tree, based on the NB-ARC domain, was inferred using the Maximum Likelihood method based on the Jones-Taylor-Thornton (JTT) model (Jones *et al*., 1992). The tree with the highest log likelihood is shown. Branch labels indicates percentage bootstrap values inferred from 1000 rapid bootstrap replicates (Stamatakis *et al*., 2008). NLRs with identical NB-ARC domains are collapsed, while for those with multiple NB-ARC domains the NB-ARC are numbered according to order in the protein. The representative sequences of the 90% sequence identity reduced redundancy set are indicated in bold. The TIR-NLR, CC-NLR, CC_R_-NLR, and CC_G10_-NLR subclades are indicated. Domain architecture is shown as in figure 3.

### Benchmarking NLR-annotation tools using RefPlantNLR

We took advantage of the RefPlantNLR dataset to benchmark NLR-annotation tools by determining their sensitivity in retrieving NLRs and accuracy in annotating NLR domain architecture. This is particularly justified because the majority of NLR prediction tools have only been evaluated using the reference Arabidopsis NLRome, which is not representative of NLR diversity across flowering plants (**Figure 1**). We selected the 5 most popular NLR annotation tools for benchmarking (**Table 1**). Since not all RefPlantNLR entries have an associated CDS entry or genomic information and some of these NLR annotation tools only take nucleotide sequences as an input we decided to proceed with the benchmarking using only the RefPlantNLR entries with CDS information (429/442). Out of the NLR-annotation tools, DRAGO2 has the highest sensitivity, retrieving all of the RefPlantNLR entries when run on amino acid sequences (**Figure 5a**, **Table 1**). NLR-Annotator has the second highest sensitivity, retrieving 421/429 (98.1%) of the sequences (**Figure 5a**, **Table 1**). It has previously been noted that NLR-Annotator does not perform well on retrieving the CC_R_-NLR subclade members (Steuernagel *et al*., 2015). Indeed, NLR-Annotator missed 6/9 (67%) of CC_R_-NLRs in the RefPlantNLR dataset, while it retrieved all TIR-NLRs and CC_G10_-NLRs, and only missed 2/309 (0.6%) of CC-NLRs (**Figure 5b**). Additionally, NLR-Annotator performs similarly on extracted genomic sequence, retrieving 371/380 (97.4%) of RefPlantNLR entries with associated genomic information (**Figure S1**). NLGenomeSweeper, which like NLR-Annotator also takes either CDS or genomic sequence as an input, performs considerably worse on genomic input as compared to CDS input retrieving 342/381 (89.8%) of RefPlantNLR entries using extracted genomic sequence as an input versus 418/429 (97.4%) of RefPlantNLR entries when CDS was used as an input (**Figure S1**). Both NLR-Annotator and NLGenomeSweeper duplicate NLRs with multiple NB-ARC domains, potentially artificially inflating the number of NLRs extracted.

**Table 1:** NLR-annotation tools

| Tool | Input | Output |  | RefPlantNLR (N = 429) |  |
| --- | --- | --- | --- | --- | --- |
|  |  | Domain annotation | NB-ARC | Sensitivity | Annotation specificity |
| DRAGO2 | AA/transcripts | Yes | No | 100%/99.3%* | 45.2%/44.9%* |
| NLGenomeSweeper | Transcripts/Genomic | Yes | Yes | 97.4%/89.8%** | 31.9%/36.5%** |
| NLR-annotator | Transcripts/Genomic | NLR-motifs | Yes | 98.1%/97.4%** | 88.1%/86.9%** |
| RGAugury | AA | Yes | No | 97.2% | 58.3% |
| RRGPredictor | AA/transcripts | Yes | No | 95.1% | 62.2% |

\*AA/CDS input
\*\*CDS/Genomic input. Gene models were available for 381 NLRs

**Figure 5:**
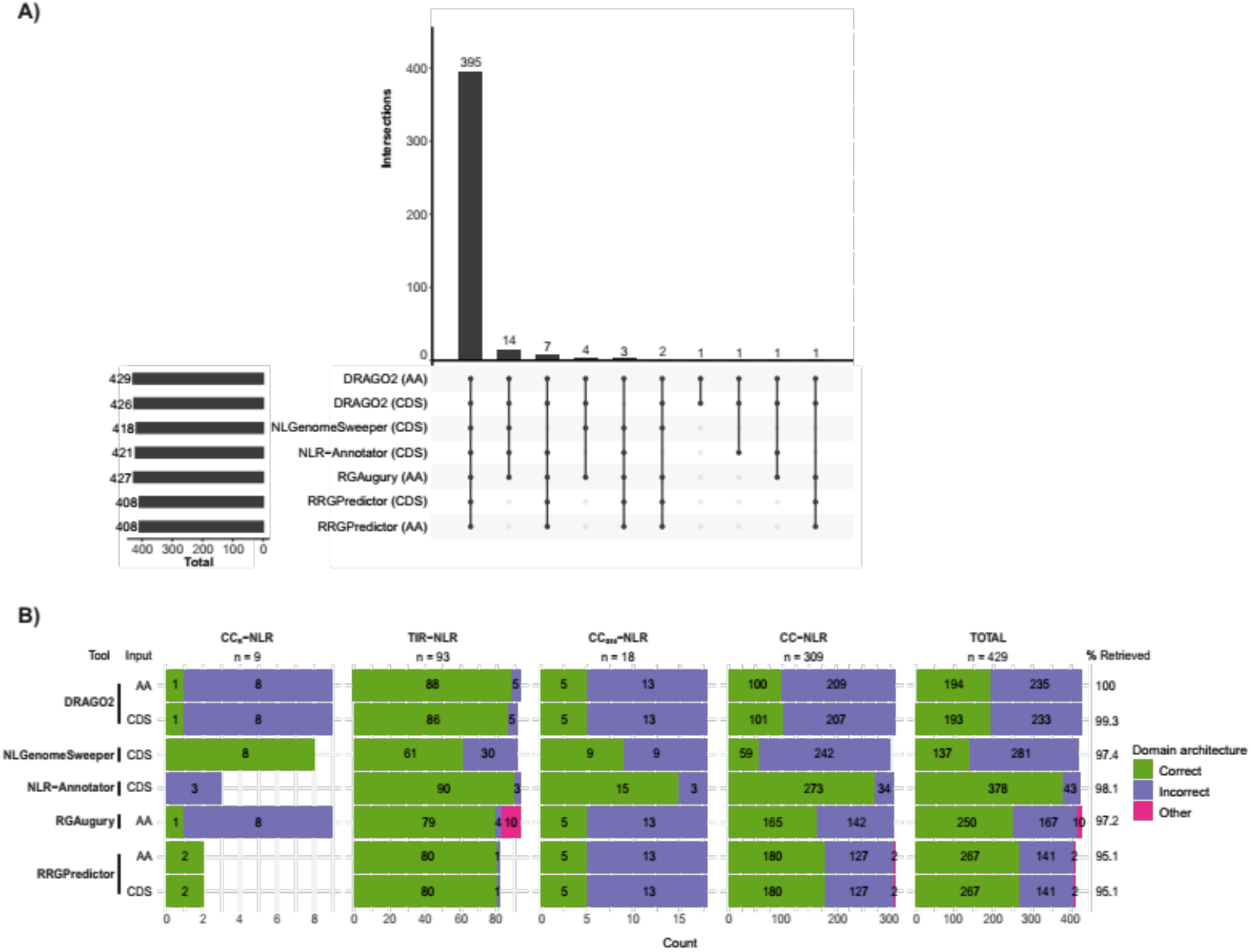
Benchmarking NLR annotation tools using RefPlantNLR. Benchmarking of NLR annotation tools using the RefPlantNLR dataset for which a CDS entry was available (N = 429). **A)** UpSet plot showing intersection of RefPlantNLR entries retrieved by each annotation tool. **B)** Domain architecture analysis produced by each NLR annotation tool per NLR subclass. Correct domain architecture is consistent with RefPlantNLR annotation, incorrect is inconsistent with RefPlantNLR annotation. Other is retrieved by NLR annotation tool but not reliably classified as NLR.

Next, we compared sensitivity and domain annotation accuracy of the NLR annotation tools according to the four main NLR subclades. While DRAGO2 is the most sensitive in retrieving NLRs, it correctly annotated the domain architecture of less than half (45%) of the RefPlantNLR sequences (**Figure 5b**, **Table 1**). Although NLR-Annotator does not automatically output a domain architecture analysis as the other tools, upon conversion of the motif analysis to domain architecture we found that NLR-Annotator has the highest domain annotation accuracy of all tools, correctly annotating 378/429 (88%) of the NLRs (**Figure 5b**, **Table 1**). The other tools did not perform much better than DRAGO2, correctly annotating between 31.9% to 62.2% of RefPlantNLR entries (**Figure 5b**, **Table 1**). When looking at the different NLR subclades it becomes clear that most tools correctly identify and annotate TIR-NLRs, while domain prediction accuracy is lower for the other NLR subclades (**Figure 5b**). The exception to this is NLR-Annotator which accurately annotates the domains of 273/309 (88%) CC-NLRs. This is possibly because NLR-Annotator was validated with the wheat genome which contains a large proportion of CC-NLRs and some of the used motifs are specific to monocot CC-NLRs (Jupe *et al*., 2012), whereas the other tools were validated with Arabidopsis, which has a higher abundance of TIR-NLRs as compared to other species. Finally, comparing these NLR-annotation tools on the reduced redundancy RefPlantNLR set revealed a similar pattern (**Figure S2**, **Appendix S1** for the full analysis). Based on the benchmarking using RefPlantNLR we find DRAGO2 to be the most sensitive tool for NLR-retrieval, while NLR-Annotator is the most sensitive tool for use on genomic input. None of the tools performs well on the domain architecture analysis except for NLR-Annotator, however, to extract such a domain architecture output from NLR-Annotator does require a substantial effort on the user side.

### NLRtracker: an NLR extraction and annotation pipeline based on the core features of RefPlantNLR

To address the limitations of the current NLR annotation tools highlighted above, we generated a novel pipeline we called NLRtracker. NLRtracker uses InterProScan (Finn *et al*., 2017) and the predefined NLR motifs (Jupe *et al*., 2012) to annotate all sequences in a given proteome or transcriptome and then extracts and annotates NLRs based on the core NLR sequence features (Late blight R1, TIR, RPW8, CC, NB-ARC, LRR, and integrated domains) found in the RefPlantNLR dataset (**Figure 6a**, **Appendix S2**). Additionally, NLRtracker extracts the NB-ARC domain for comparative phylogenetic analysis.

**Figure 6:**
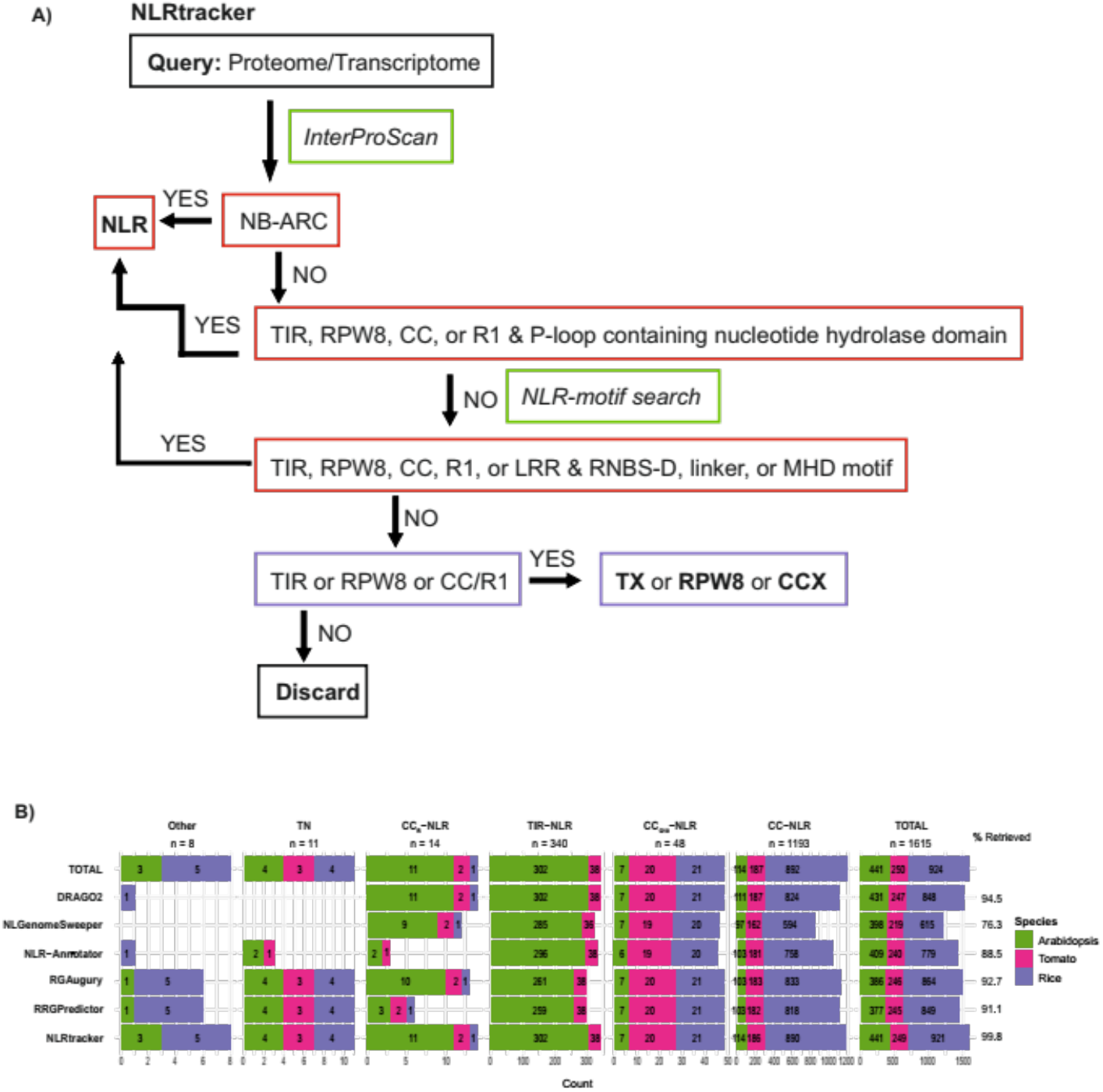
Sensitivity of NLRtracker compared to other annotation tools using Arabidopsis, tomato, and rice RefSeq genomes. Benchmarking of NLR annotation tools using the Arabidopsis, rice and tomato RefSeq genomes. **A)** NLRtracker pipeline. InterProScan and predefined NLR-motifs are used to group sequences into different categories. **B)** Number of NLRs retrieved in each NLR subclass per species.

We compared NLRtracker to the other NLR-annotation tools on the Arabidopsis, tomato, and rice RefSeq genomes. In this way we could also assess the accuracy of each NLR-annotation tool, something which is not possible with the RefPlantNLR dataset. Using all tools, we extracted a total of 1615 NLRs from the reference Arabidopsis (N = 441), tomato (N = 250), and rice (N = 924) genomes (**Figure 6b**). The total number of NLRs belonging to each subclade in each species is reflected in the distribution of each subclade in these species in the RefPlantNLR dataset (**Figure 6b**). In addition to the four main subclades of NLRs, we also retrieved a highly conserved TIR-NB-ARC (TN) class of proteins which phylogenetically clusters separately from all other plant NLRs and whose gene structure is clearly distinct from other TIR-NLRs (Meyers *et al*., 2002), as well as certain NB-ARC containing proteins which did not clearly belong to any of these clades. This included an Arabidopsis NB-ARC protein with an integrated ZBTB8B domain in between the NB-ARC and the LRR, as well as a rice protein containing an NB-ARC domain with a C-terminal armadillo-type SSFR (**Figure 6b, Appendix S3** for the full analysis).

Using this dataset, we evaluated the sensitivity and specificity of each NLR-annotation tool. Sensitivity was defined as the total percentage NLRs retrieved out of the total NLR dataset, while specificity was defined as the total number of sequences annotated as NLRs being genuine NLRs. False positives could include TIR- or RPW8-only proteins annotated as genuine NLRs, or unrelated sequences annotated as NLRs. On this dataset, NLRtracker had the highest sensitivity (retrieving 1611/1615 (99.8%) NLRs with 100% accuracy (**Table 2**, **Figure 6b**, **Figure S3**). The four missed sequences were retrieved by DRAGO2 and included one NLR with an N-terminal CC-domain and a C-terminal LRR, but which did not get annotated with an NB-ARC domain using InterProScan or the predefined NLR motifs, and three truncated proteins, two of which contain an LRR domain while one does not get annotated with any domain using InterProScan.

**Table 2:**
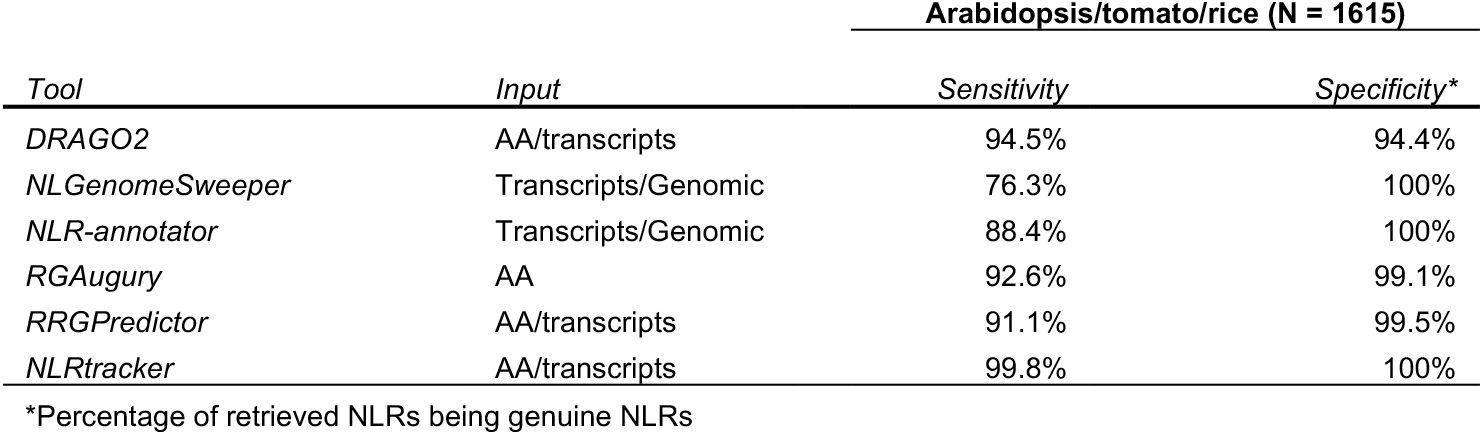
Extraction of NLRs from the Arabidopsis, tomato, and rice RefSeq proteomes Arabidopsis/tomato/rice (N = 1615)

Of the pre-existing tools DRAGO2 was the most sensitive, retrieving 1526/1615 (94.5%) NLRs, however, it also was the least accurate method extracting 91 false positives (**Table 1**, **Figure S3**). These false positives were predominantly proteins containing a P-loop containing nucleoside triphosphate hydrolase domain unrelated to the NB-ARC domain. Similarly, RGAugury extracted 13 such false positives. By contrast, the 8 false positives extracted by RRGPredictor are RPW8-containing proteins lacking an NB-ARC domain. In conclusion, the NLRtracker tool we developed here is more sensitive and more accurate than previously available tools for extracting NLRs from a given plant proteome/transcriptome. Additionally, NLRtracker facilitates domain architecture analysis and phylogenetic analysis. Combining the extracted NB-ARC domain generated by NLRtracker with the RefPlantNLR extracted NB-ARC dataset (**Supplemental dataset 10**) should greatly facilitate comparative phylogenetics and reveal the phylogenetic relationships of a newly annotated NLR. Nevertheless, the quality of the output remains dependent on the quality of the input sequences, and none of these tools is able to determine whether an extracted sequence represents a genuine NLR, as in having a genuine NB-ARC domain or consisting of a full-length protein. For example, 94/1615 extracted proteins do not get annotated with an NB-ARC domain, of which 44 do not get annotated with a P-loop containing nucleoside triphosphate hydrolases domain but do contain NB-ARC-specific motifs in combination with domains found in NLRs. Some of these may represent genuine NLRs such as Pb1 or RXL, that have undergone regressive evolution, whereas other may be partial or pseudogenes.

### Additional applications of the RefPlantNLR dataset

We showed that RefPlantNLR is useful for benchmarking and improving NLR annotation tools. Additional uses of the dataset include providing reference points for newly discovered NLRs with NLRtracker feeding into the large-scale phylogenetic analyses that are necessary for classifying NLRomes. Phylogenetic analyses would help assign NLRs to subclades and provide a basis for generating hypotheses about the function and mode of action of novel NLRs which phylogenetically cluster with experimentally validated NLRs. This type of phylogenetic information can be combined with other features such as genetic clustering and has, for instance, proven valuable in previous work on rice and solanaceous NLRs (Białas *et al*., 2017; Wu *et al*., 2017) and for defining the CC_G10_-NLR class (Lee *et al*., 2020).

Furthermore, known mutants and sequence variants can be mapped onto a phylogenetic framework, such as the RefPlantNLR tree (**Figure 4**). For example, the CC-NLR ZAR1 and TIR-NLR ROQ1 are bound to ATP in their activated form (Wang *et al*., 2019; Martin *et al*., 2020), whereas the TIR-NLR RPP1 is bound to ADP in its activated form (Ma *et al*., 2020). RefPlantNLR has already proven useful in interpreting a feature of the recently elucidated structure of the RPP1 resistosome (Ma *et al*., 2020). The authors used RefPlantNLR to determine that although most CC-NLRs contain a TT/SR motif in which the arginine interacts with ATP, a subset of TIR-NLRs contain a charged or polar substitution creating a TTE/Q motif interacting with ADP in the activated form (Ma *et al*., 2020). Interestingly a phylogenetically distinct subgroup of CC-NLRs known as the MIC1 group (Bailey *et al*., 2018) is an exception to this rule by having a TTE/Q motif in their ADP binding pocket and thus may also retain ADP binding when activated. This example shows how a carefully curated reference dataset like RefPlantNLR can facilitate data interpretation and hypothesis generation.

RefPlantNLR highlights the under-studied plant species of NLR biology. **Table S4** reveals that ~80% (48 out of 59) of the major seed plant clades recently defined by Smith and Brown (Smith and Brown, 2018) do not have a single experimentally validated NLR. Certain taxa have subfamily-specific contractions and expansions, and hence may contain unexplored genetic and biochemical diversity of NLR function. Looking forward, combining the output of NLRtracker with RefPlantNLR may highlight understudied subgroups of NLRs even within well-studied organisms. For example, while all currently studied CC_R_-NLRs act as helper NLRs for TIR-NLRs, it has been reported that the CC_R_-NLR subfamily has experienced clade-specific expansions in gymnosperms and rosids, pointing to potential biochemical specialization of this subfamily in these taxa (Van Ghelder *et al*., 2019). In addition, although NLRs have been reported in non-seed plants and some of these appear to have distinct N-terminal domains (Andolfo *et al*., 2019), their experimental validation is still lacking.

The RefPlantNLR dataset has inherent limitations due to its focus on experimentally validated NLRs. First, it is biased towards a few well studied model species and crops as illustrated in **Figure 1**. Additionally, RefPlantNLR entries are somewhat redundant with particular NLR allelic series, such as the monocot MLA and spinach alpha-WOLF, being overrepresented in the dataset (**Figure 1**, **Figure 4**). These issues, notably redundancy, will need to be considered for certain applications where it may be preferable to use the reduced redundancy dataset (**Supplemental dataset 13, 14**).

### Conclusion

We hope that the RefPlantNLR resource will contribute to moving the field beyond a uniform view of NLR structure and function. It is now evident that NLRs turned out to be more structurally and functionally diverse than anticipated. Whereas a number of plant NLRs have retained the presumably ancestral three domain architecture of the TIR/CC_R_/CC_G10_/CC fused to the NB-ARC and LRR domains, many NLRs have diversified into specialized proteins with degenerated features and extraneous non-canonical integrated domains (Sarris *et al*., 2016; Kroj *et al*., 2016; Adachi, Derevnina, *et al*., 2019). Therefore, it is time to question holistic concepts such as effector-triggered immunity (ETI) and appreciate the wide structural and functional diversity of NLR-mediated immunity. More specifically, a robust phylogenetic framework of plant NLRs should be fully integrated into the mechanistic study of these exceptionally diverse proteins.

## MATERIAL & METHODS

### Sequence retrieval

RefPlantNLR was assembled by manually crawling the literature for experimentally validated NLRs according to the criteria described in the results section. NLRs are defined as having a NB-ARC and at least one additional domain. Where possible the amino acid and nucleotide sequences were taken from GenBank. For some NLRs, only the mRNA has been deposited and no genomic locus information was present. When GenBank records were not available, the sequences were extracted from the matching whole-genome sequences projects or from articles and patents describing the identification of these NLRs.

### Domain annotation

Protein sequences were annotated with CATH-Gene3D (v4.2.0) (Dawson *et al*., 2017), SUPERFAMILY (v1.75) (Gough *et al*., 2001), PRINTS (v42.0) (Attwood *et al*., 2000), PROSITE profiles (v2019_11) (Sigrist *et al*., 2013), SMART (v7.1) (Letunic *et al*., 2015), CDD (v3.17) (Marchler-Bauer *et al*., 2015), and Pfam (v33.1) (El-Gebali *et al*., 2019) identifiers using InterProScan (v5.47-82.0) (Finn *et al*., 2017) and predefined NLR-motifs (Jupe *et al*., 2012) using the meme-suite (v5.1.1) (Bailey *et al*., 2009). A custom R script (**Appendix S4**) was used to convert the InterProScan output to the final GFF3 annotation and extract the NB-ARC domain. We routinely use Geneious Prime (v2020.2.4) (https://www.geneious.com) to visualize these annotations on the sequence. The NLR-associated signature motifs/domain IDs are:

- Late blight resistance protein R1: PF12061
- Rx-type CC: PF18052, cd14798
- RPW8-type CC: PF05659, PS51153
- TIR: PF01582, PF13676, G3DSA:3.40.50.10140, SSF52200, PS50104, SM00255
- NB-ARC: PF00931, G3DSA:1.10.8.430
- NB-ARC used for phylogenetic analysis: overlap of G3DSA:3.40.50.300, SSF52540, G3DSA:1.10.8.430, and PF00931 domains
- NB-ARC associated motifs: motif 2, 7, and 8 from Jupe *et al*., (2012) corresponding to the CC_R_/CC_G10_/CC-type RNBS-D, MHD, and linker motifs of the NB-ARC domain, respectively
- LRRs: G3DSA:3.80.10.10, PF08263, PF07723, PF07725, PF12799, PF13306, PF00560, PF13516, PF13855, SSF52047, SSF52058, SM00367, SM00368, SM00369, PF18837, PF01463, SM00082, SM00013, PF01462, PF18831, and PF18805
- Other: any other Pfam, SUPERFAMILY, and/or CATH-Gene3D annotation. Additionally, we included the PROSITE Profiles signatures PS51697 (ALOG domain) and PS50808 (zinc-finger BED domain), and the SMART signature SM00614 (zinc-finger BED domain).

### Sequence deduplication

The NLR amino acid sequences were clustered using CD-HIT at 90% sequence identity (v4.8.1; Fu *et al*., 2012; Usage: cd-hit -i RefPlantNLR.fasta -o RefPlantNLR _90 -c 0.90 -n 5 -M 16000 -d 0). A custom R script (**Appendix S4**) was used to assign representative sequences per cluster per genus, i.e. if a single cluster contained sequences from multiple genera we assigned a representative sequence per genus. The reduced redundancy sequences are provided in **Supplemental dataset 13, 14**.

### Phylogenetics

The NB-ARC domain of all NLRs were extracted and deduplicated using a custom R script (**Appendix S4**). For sequences containing multiple NB-ARC domains the extracted NB-ARC domain was numbered according to occurrence in the protein. Sequences were aligned using Clustal Omega (Sievers *et al*., 2011), and all positions with less than 95% site coverage were removed using QKphylogeny (Moscou, 2020) (**Supplemental dataset 11**). RAxML (v8.2.12) (Stamatakis, 2014) was used (usage: raxmlHPC-PTHREADS-AVX -T 6 -s RefPlantNLR.phy -n RefPlantNLR -m PROTGAMMAAUTO -f a -# 1000 -x 8153044963028367 -p 644124967711489) to infer the evolutionary history using the Maximum Likelihood method based on the JTT model (Jones *et al*., 1992). Bootstrap values from 1000 rapid bootstrap replicates as implemented in RAxML are shown (Stamatakis *et al*., 2008) (**Supplemental dataset 12**). The RefPlantNLR phylogeny was rooted on the TIR-/CC_R_-subclade and edited using the iTOL suite (v5.5.1; Letunic and Bork, 2019).

### Figures describing RefPlantNLR

The figures describing the RefPlantNLR dataset were generated using a custom R script (**Appendix S5**).

### Benchmarking RefPlantNLR

For benchmarking using the RefPlantNLR dataset we used DRAGO2 (DRAGO2-API, Osuna-Cruz *et al*., 2018), NLGenomeSweeper (v1.2.0, Toda *et al*., 2020; dependencies: Python 3.8, NCBI-BLAST+ (v2.11.0+), MUSCLE aligner (v3.8.1551), SAMtools (v1.9-50-g18be38a), bedtools (v2.27.1-9-g5f83cacb), HMMER (v3.3.1), InterProScan (v5.47-82.0), TransDecoder (v5.5.0)), NLR-Annotator (Steuernagel *et al*., 2020; dependencies: meme-suite (v5.1.1), NLR-Parser (v3;Steuernagel *et al*., 2015), Oracle Java SE Development Kit 11.0.9), RGAugury (Li *et al*., 2016; dependencies: CViT, HMMER, InterProScan, ncoils, NCBI-BLAST+, Pfamscan, Phobius), and RRGPredictor (Santana Silva and Micheli, 2020; dependencies: InterProScan) using either amino acid, CDS, and/or the extracted NLR genomic loci as an input. Since NLGenomeSweeper and NLR-Annotator only accept nucleotide input, while RGAugury only accepts amino acid input, we only used RefPlantNLR entries for which CDS was available in the direct comparison. For the domain analysis only the TIR, RxN-type CC, RPW8-type CC, NB-ARC, and LRR domains were considered. Additionally, sequentially duplicated domains were compressed in a single annotation. A custom R script was used to generate the analysis (**Appendix S1**).

### Description of NLRtracker

NLRtracker (**Appendix S2**) runs InterProScan (v5.47-82.0) (Finn *et al*., 2017) and FIMO from the meme-suite (v5.1.1) (Bailey *et al*., 2009) using predefined NLR-motifs (Jupe *et al*., 2012). An R script which depends on the Tidyverse (Wickham *et al*., 2019) extracts sequences containing NLR-associated domains and classifies them into different subgroups:

- NLR: containing an NB-ARC domain
- Degenerate NLR: containing RxN-type CC, late blight resistance protein R1, RPW8-type CC, or TIR in combination with a P-loop containing nucleotide hydrolase domain not overlapping with other annotations or containing a RxN-type CC, late blight resistance protein R1, RPW8-type CC, TIR, or LRR with a RNBS-D, linker, and/or MHD motif
- TX: TIR domain containing protein lacking a P-loop containing nucleotide hydrolase domain and RNBS-D, linker, and/or MHD motif
- CCX: RxN-type CC or late blight resistance protein R1 domain containing protein lacking a P-loop containing nucleotide hydrolase domain and RNBS-D, linker, and/or MHD motif
- RPW8: RPW8-type CC domain containing protein lacking a P-loop containing nucleotide hydrolase domain and RNBS-D, linker, and/or MHD motif
- MLKL: containing HeLo domain of plant-specific mixed-lineage kinase domain like proteins (PF06760; DUF1221) (Mahdi *et al*., 2020)

NLRtracker outputs the domain architecture analysis, as well as the domain boundaries. Additionally, the NB-ARC is extracted facilitating phylogenetic analysis. The current version of NLRtracker can be accessed through GitHub (https://github.com/slt666666/NLRtracker).

### Benchmarking on Arabidopsis, tomato and rice genomes

The NCBI RefSeq proteomes of Arabidopsis (*Arabidopsis thaliana* ecotype Col-0; genome assembly GCF_000001735.4; TAIR and Araport annotation), tomato (*Solanum lycopersicum* cv. Heinz 1706; genome assembly GCF_000188115.4; RefSeq annotation v103), rice (*Oryza sativa* group *Japonica* cv. Nipponbare; genome assembly GCF_001433935.1; RefSeq annotation v102) were downloaded from NCBI. We used NLRtracker, DRAGO2, RGAugury, and RRGPredictor on amino acid sequences, while we used the extracted CDS from the genomic sequence as an input for NLGenomeSweeper and NLR-Annotator.

NLRs were grouped in different subclades based on phylogenetic clustering with the RefPlantNLR CC_R_-NLR, TIR-NLR, CC_G10_-NLR, and CC-NLR subgroups, while those that did not clearly fall into any of these groups but contained a TIR-domain and P-loop containing nucleotide hydrolase domain were classified as TN subclade members. The remainder was grouped together and classified as other. Proteins which were extracted but did not belong to the NLR subfamily were manually inspected and classified as false positives. Additionally, TIR- or RPW8-only (TX and RPW8 respectively) proteins extracted as NLRs were marked as false positives. A custom R script was used to generate the analysis (**Appendix S3**).

## Data availability and updates

Up to date versions of RefPlantNLR can be accessed via Zenodo at http://doi.org/10.5281/zenodo.3936022. This project is part of the OpenPlantNLR community on Zenodo: https://zenodo.org/communities/openplantnlr

## ACKNOWLEDGEMENTS

We thank Adeline Harant, Philip Carella, and Hiral Shah for useful comments and feedback, and Aleksandra Białas for the domain architecture illustrations.

## FUNDING

This work has been supported by the Gatsby Charitable Foundation, Biotechnology and Biological Sciences Research Council (BBSRC), European Research Council (ERC) and BASF Plant Science. The funders had no role in study design, data collection and analysis, decision to publish, or preparation of the manuscript.

## COMPETING INTERESTS

The authors receive funding from industry on NLR biology.

## AUTHOR CONTRIBUTIONS

Conceptualization: JK, SK, TS; Data curation: JK; Formal analysis: JK, TS, HA; Investigation: JK, TS, HA; Code: TS, JK; Supervision: SK; Funding acquisition: SK; Project administration: SK; Writing initial draft: JK; Editing: JK, TS, SK.

## SUPPLEMENTAL DATA

**Table S1: Description of RefPlantNLR.**

**Table S2: Description of animal, bacterial, and archaeal NB-ARC-domain containing proteins.**

**Table S3: Description of NLR-associated proteins.**

**Table S4: Plant orders represented in RefPlantNLR.**

**Supplemental dataset 1: Amino acid sequences of RefPlantNLR entries (fasta format).** This file contains 442 amino acid sequences.

**Supplemental dataset 2: CDS sequences of RefPlantNLR entries (fasta format).** This file contains 427 CDS sequences. For 15 RefPlantNLR entries no CDS sequence could be retrieved.

**Supplemental dataset 3: Annotated genomic sequences of RefPlantNLR entries (GenBank flat file format).** This file contains 354 genomic loci containing the gene models of 387 RefPlantNLR entries.

**Supplemental dataset 4: Amino acid sequences of animal, bacterial, and archaeal NB-ARC domain containing proteins (fasta format).** This file contains 13 amino acid sequences.

**Supplemental dataset 5: Amino acid sequences of NLR-associated entries (fasta format).** This file contains 15 amino acid sequences.

**Supplemental dataset 6: CDS sequences of NLR-associated entries (fasta format).** This file contains 15 CDS sequences. For 1 entry no CDS sequence could be retrieved.

**Supplemental dataset 7: Annotated genomic sequences of NLR-associated entries (GenBank flat file format).** This file contains 13 genomic loci containing the gene models of 14 NLR-associated entries.

**Supplemental dataset 8: Domain and motif annotation of the RefPlantNLR amino acid sequences, animal, bacterial, and archaeal NB-ARC domain containing proteins, and NLR-associated proteins (GFF3 format).** This file contains the InterProScan annotation of 470 amino acid sequences.

**Supplemental dataset 9: Amino acid sequences of the extracted RefPlantNLR NB-ARC domains (fasta format).** This file contains 450 NB-ARC domain amino acid sequences belonging to 442 RefPlantNLR entries.

**Supplemental dataset 10: Amino acid sequences of the unique RefPlantNLR extracted NB-ARC domains (fasta format).** This file contains 370 unique NB-ARC domains amino acid sequences.

**Supplemental dataset 11: Clustal Omega alignment of the unique RefPlantNLR extracted NB-ARC domains (PHYLIP format).** This file contains the Clustal Omega alignment of 369 unique NB-ARC domains with all positions with less than 95% coverage removed. Pb1 and RXL were omitted from this alignment.

**Supplemental dataset 12: NB-ARC domain phylogeny of the RefPlantNLR entries using the Maximum likelihood method (Newick format).** This file contains the phylogenetic analysis of the NB-ARC domain of the RefPlantNLR entries using the JTT method.

**Supplemental dataset 13: Amino acid sequences of the non-redundant RefPlantNLR entries (fasta format).** This file contains 285 amino acid sequences representing the non-redundant RefPlantNLR entries at a 90% identity threshold per genus.

**Supplemental dataset 14: Amino acid sequences of the non-redundant RefPlantNLR entries (fasta format).** This file contains 281 amino acid sequences representing the unique non-redundant RefPlantNLR at a 90% identity threshold per genus extracted NB-ARC domains.

**Appendix S1: Benchmarking using RefPlantNLR. Scripts and data.**

**Appendix S2: NLRtracker. Scripts.**

**Appendix S3: Benchmarking using Arabidopsis, tomato, and rice proteomes. Scripts and data.**

**Appendix S4: Scripts and data to convert annotations, extract NB-ARC domain and assign representative entries.**

**Appendix S5: R script used to generate figures describing RefPlantNLR.**

---

**Figure S1:**
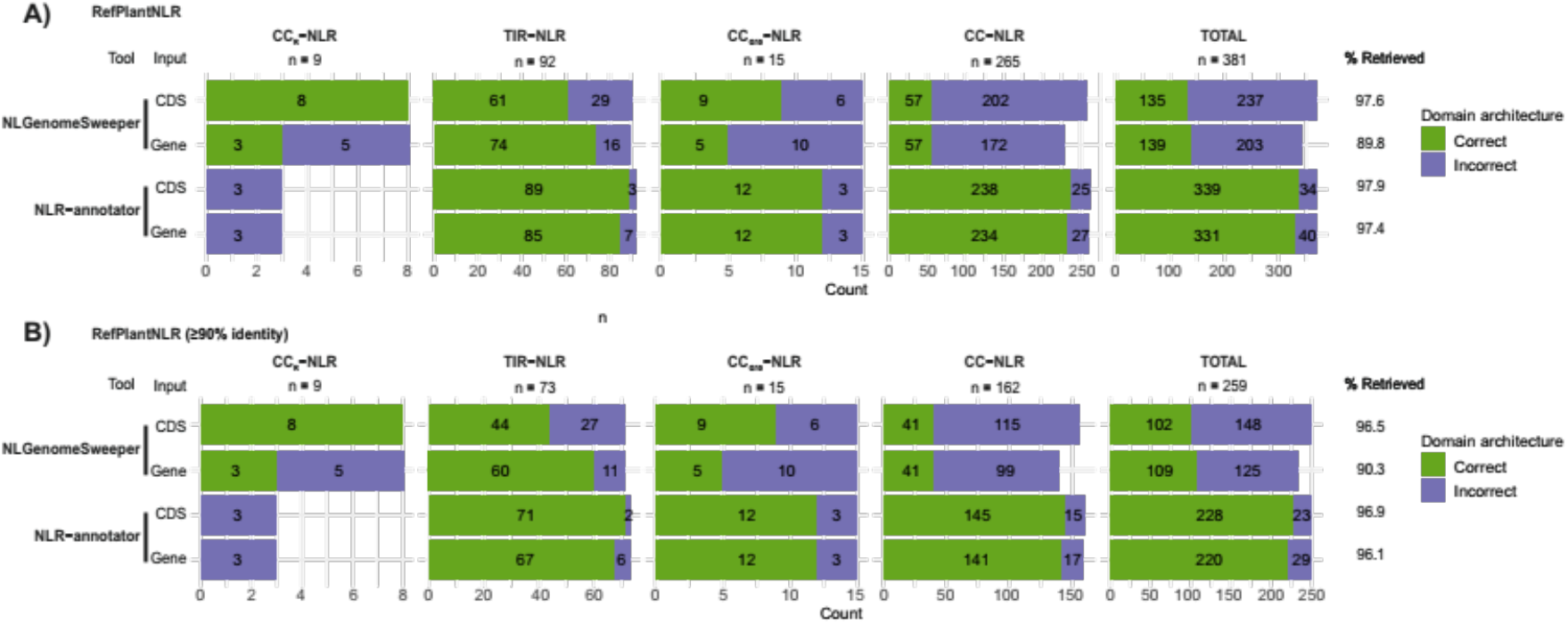
Comparison of NLR-Annotator and NLGenomeSweeper on CDS versus genomic input. NLR-Annotator and NLGenomeSweeper were run on CDS or genomic input. **A)** Domain architecture analysis of NLR-Annotator and NLGenomeSweeper run on CDS or genomic input from each RefPlantNLR entry. Only entries for which a genomic locus was available were considered (N = 381). **B)** Same as **A)** for the representative dataset (N = 259). Correct domain architecture is consistent with RefPlantNLR annotation, incorrect is inconsistent with RefPlantNLR annotation.

**Figure S2:**
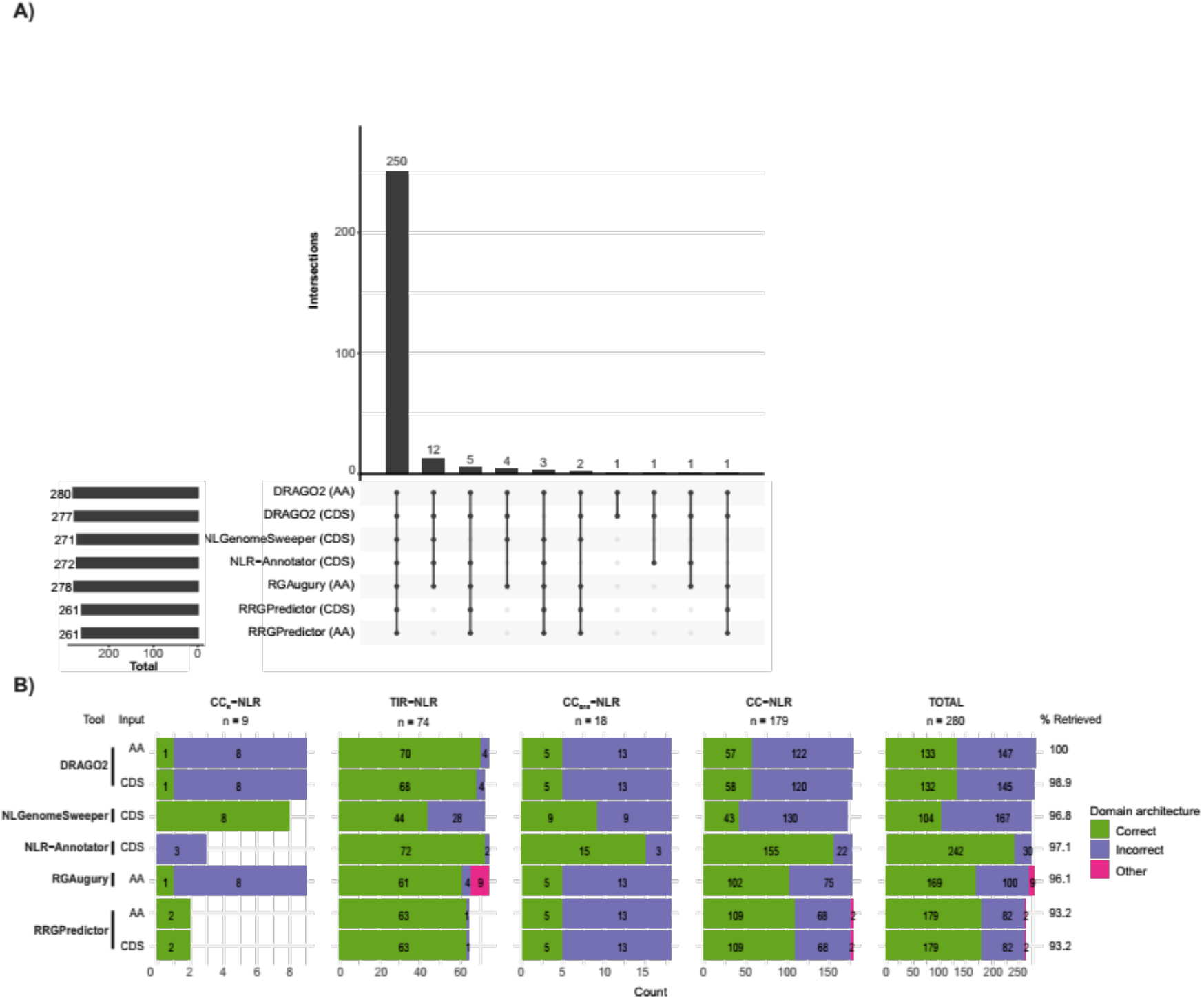
Benchmarking NLR annotation tools using reduced redundancy RefPlantNLR entries. Benchmarking of NLR annotation tools using the reduced redundancy RefPlantNLR dataset for which a CDS entry was available (N = 280). **A)** UpSet plot showing intersection of RefPlantNLR entries retrieved by each annotation tool. **B)** Domain architecture analysis produced by each NLR annotation tool per NLR subclass. Correct domain architecture is consistent with RefPlantNLR annotation, incorrect is inconsistent with RefPlantNLR annotation. Other is retrieved by NLR annotation tool but not reliably classified as NLR.

**Figure S3:**
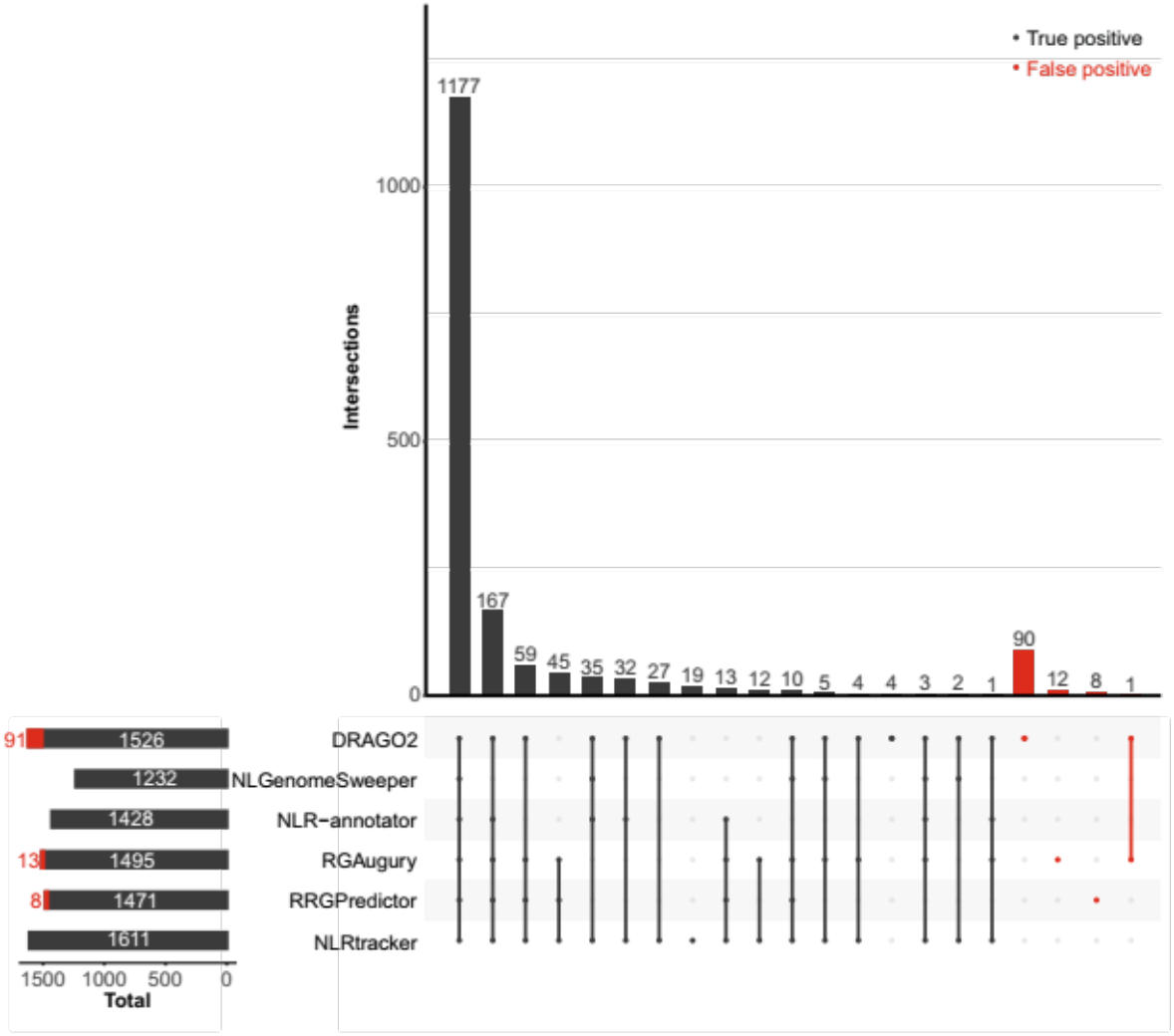
Sensitivity and accuracy of NLRtracker compared to other annotation tools using Arabidopsis, tomato, and rice RefSeq genomes. Benchmarking of NLR annotation tools using the Arabidopsis, rice and tomato RefSeq genomes. UpSet plot showing intersection of NLRs retrieved by each annotation tool. False positive annotations are marked in red.

## Notes

http://doi.org/10.5281/zenodo.3936022

### Summary of Updates

This updated version of RefPlantNLR now consists of 442 NLRs from 31 genera belonging to 11 orders of flowering plants. We also used RefPlantNLR to benchmark the five most popular NLR annotation tools. Guided by this analysis, we developed a new pipeline, NLRtracker, which extracts and annotates NLRs based on the core features found in the RefPlantNLR dataset.

## REFERENCES

Adachi, H., Contreras, M.P., Harant, A., et al. (2019) An N-terminal motif in NLR immune receptors is functionally conserved across distantly related plant species. eLife, 8, e49956.

Adachi, H., Derevnina, L. and Kamoun, S. (2019) NLR singletons, pairs, and networks: evolution, assembly, and regulation of the intracellular immunoreceptor circuitry of plants. Curr. Opin. Plant Biol., 50, 121–131.

Andolfo, G., Di Donato, A., Chiaiese, P., De Natale, A., Pollio, A., Jones, J.D.G., Frusciante, L. and Ercolano, M.R. (2019) Alien domains shaped the modular structure of plant NLR proteins. Genome Biol. Evol., 11, 3466–3477.

Araújo, A.C. de, Fonseca, F.C.D.A., Cotta, M.G., Alves, G.S.C. and Miller, R.N.G. (2020) Plant NLR receptor proteins and their potential in the development of durable genetic resistance to biotic stresses. Biotechnol. Res. Innov.

Attwood, T.K., Croning, M.D.R., Flower, D.R., Lewis, A.P., Mabey, J.E., Scordis, P., Selley, J.N. and Wright, W. (2000) PRINTS-S: the database formerly known as PRINTS. Nucleic Acids Res., 28, 225–227.

Bailey, P.C., Schudoma, C., Jackson, W., Baggs, E., Dagdas, G., Haerty, W., Moscou, M. and Krasileva, K.V. (2018) Dominant integration locus drives continuous diversification of plant immune receptors with exogenous domain fusions. Genome Biol., 19, 23.

Bailey, T.L., Boden, M., Buske, F.A., Frith, M., Grant, C.E., Clementi, L., Ren, J., Li, W.W. and Noble, W.S. (2009) MEME Suite: tools for motif discovery and searching. Nucleic Acids Res., 37, W202–W208.

Barragan, C.A., Wu, R., Kim, S.-T., et al. (2019) RPW8/HR repeats control NLR activation in Arabidopsis thaliana. PLOS Genet., 15, e1008313.

Białas, A., Zess, E.K., De la Concepcion, J.C., et al. (2017) Lessons in effector and NLR biology of plant-microbe systems. Mol. Plant. Microbe Interact., 31, 34–45.

Biezen, E.A. van der and Jones, J.D.G. (1998) Plant disease-resistance proteins and the gene-for-gene concept. Trends Biochem. Sci., 23, 454–456.

Cesari, S., Bernoux, M., Moncuquet, P., Kroj, T. and Dodds, P.N. (2014) A novel conserved mechanism for plant NLR protein pairs: the “integrated decoy” hypothesis. Plant-Microbe Interact., 5, 606.

Chae, E., Bomblies, K., Kim, S.-T., et al. (2014) Species-wide genetic incompatibility analysis identifies immune genes as hot spots of deleterious epistasis. Cell, 159, 1341–1351.

Dangl, J.L., Horvath, D.M. and Staskawicz, B.J. (2013) Pivoting the plant immune system from dissection to deployment. Science, 341, 746–751.

Dawson, N.L., Lewis, T.E., Das, S., Lees, J.G., Lee, D., Ashford, P., Orengo, C.A. and Sillitoe, I. (2017) CATH: an expanded resource to predict protein function through structure and sequence. Nucleic Acids Res., 45, D289–D295.

Dyrka, W., Coustou, V., Daskalov, A., et al. (2020) Identification of NLR-associated amyloid signaling motifs in filamentous bacteria. bioRxiv, 2020.01.06.895854.

El-Gebali, S., Mistry, J., Bateman, A., et al. (2019) The Pfam protein families database in 2019. Nucleic Acids Res., 47, D427–D432.

Finn, R.D., Attwood, T.K., Babbitt, P.C., et al. (2017) InterPro in 2017—beyond protein family and domain annotations. Nucleic Acids Res., 45, D190–D199.

Fu, L., Niu, B., Zhu, Z., Wu, S. and Li, W. (2012) CD-HIT: accelerated for clustering the next-generation sequencing data. Bioinformatics, 28, 3150–3152.

Gao, L., Altae-Tran, H., Böhning, F., et al. (2020) Diverse enzymatic activities mediate antiviral immunity in prokaryotes. Science, 369, 1077–1084.

Gough, J., Karplus, K., Hughey, R. and Chothia, C. (2001) Assignment of homology to genome sequences using a library of hidden Markov models that represent all proteins of known structure. J. Mol. Biol., 313, 903–919.

Hayashi, N., Inoue, H., Kato, T., et al. (2010) Durable panicle blast-resistance gene *Pb1* encodes an atypical CC-NBS-LRR protein and was generated by acquiring a promoter through local genome duplication. Plant J., 64, 498–510.

Hoorn, R.A.L. van der and Kamoun, S. (2008) From guard to decoy: a new model for perception of plant pathogen effectors. Plant Cell Online, 20, 2009–2017.

Jacob, F., Vernaldi, S. and Maekawa, T. (2013) Evolution and Conservation of Plant NLR Functions. Front. Immunol., 4.

Jones, D.T., Taylor, W.R. and Thornton, J.M. (1992) The rapid generation of mutation data matrices from protein sequences. Bioinformatics, 8, 275–282.

Jones, J.D.G., Vance, R.E. and Dangl, J.L. (2016) Intracellular innate immune surveillance devices in plants and animals. Science, 354.

Jupe, F., Pritchard, L., Etherington, G.J., et al. (2012) Identification and localisation of the NB-LRR gene family within the potato genome. BMC Genomics, 13, 75.

Kourelis, J. and Hoorn, R.A.L. van der (2018) Defended to the nines: 25 years of resistance gene cloning identifies nine mechanisms for R protein function. Plant Cell, 30, 285–299.

Kroj, T., Chanclud, E., Michel-Romiti, C., Grand, X. and Morel, J.-B. (2016) Integration of decoy domains derived from protein targets of pathogen effectors into plant immune receptors is widespread. New Phytol., 210, 618–626.

Lee, H.-Y., Mang, H., Choi, E., Seo, Y.-E., Kim, M.-S., Oh, S., Kim, S.-B. and Choi, D. (2020) Genome-wide functional analysis of hot pepper immune receptors reveals an autonomous NLR clade in seed plants. New Phytol.

Letunic, I. and Bork, P. (2019) Interactive Tree Of Life (iTOL) v4: recent updates and new developments. Nucleic Acids Res., 47, W256–W259.

Letunic, I., Doerks, T. and Bork, P. (2015) SMART: recent updates, new developments and status in 2015. Nucleic Acids Res., 43, D257–D260.

Li, P., Quan, X., Jia, G., Xiao, J., Cloutier, S. and You, F.M. (2016) RGAugury: a pipeline for genome-wide prediction of resistance gene analogs (RGAs) in plants. BMC Genomics, 17, 852.

Ma, S., Lapin, D., Liu, L., et al. (2020) Direct pathogen-induced assembly of an NLR immune receptor complex to form a holoenzyme. Science, 370.

Mahdi, L.K., Huang, M., Zhang, X., et al. (2020) Discovery of a family of mixed lineage kinase domain-like proteins in plants and their role in innate immune signaling. Cell Host Microbe.

Marchal, C., Zhang, J., Zhang, P., et al. (2018) BED-domain-containing immune receptors confer diverse resistance spectra to yellow rust. Nat. Plants, 4, 662–668.

Marchler-Bauer, A., Derbyshire, M.K., Gonzales, N.R., et al. (2015) CDD: NCBI’s conserved domain database. Nucleic Acids Res., 43, D222–D226.

Martin, R., Qi, T., Zhang, H., Liu, F., King, M., Toth, C., Nogales, E. and Staskawicz, B.J. (2020) Structure of the activated ROQ1 resistosome directly recognizing the pathogen effector XopQ. Science, 370.

Meyers, B.C., Morgante, M. and Michelmore, R.W. (2002) TIR-X and TIR-NBS proteins: two new families related to disease resistance TIR-NBS-LRR proteins encoded in *Arabidopsis* and other plant genomes. Plant J., 32, 77–92.

Moscou, M. (2020) matthewmoscou/QKphylogeny, Available at: https://github.com/matthewmoscou/QKphylogeny.

Osuna-Cruz, C.M., Paytuvi-Gallart, A., Di Donato, A., Sundesha, V., Andolfo, G., Aiese Cigliano, R., Sanseverino, W. and Ercolano, M.R. (2018) PRGdb 3.0: a comprehensive platform for prediction and analysis of plant disease resistance genes. Nucleic Acids Res., 46, D1197–D1201.

Santana Silva, R.J. and Micheli, F. (2020) RRGPredictor, a set-theory-based tool for predicting pathogen-associated molecular pattern receptors (PRRs) and resistance (R) proteins from plants. Genomics, 112, 2666–2676.

Sarris, P.F., Cevik, V., Dagdas, G., Jones, J.D.G. and Krasileva, K.V. (2016) Comparative analysis of plant immune receptor architectures uncovers host proteins likely targeted by pathogens. BMC Biol., 14, 8.

Schaafsma, G.C.P. and Vihinen, M. (2018) Representativeness of variation benchmark datasets. BMC Bioinformatics, 19, 461.

Seong, K., Seo, E., Witek, K., Li, M. and Staskawicz, B. (2020) Evolution of NLR resistance genes with noncanonical N-terminal domains in wild tomato species. New Phytol.

Shao, Z.-Q., Xue, J.-Y., Wu, P., Zhang, Y.-M., Wu, Y., Hang, Y.-Y., Wang, B. and Chen, J.-Q. (2016) Large-scale analyses of angiosperm nucleotide-binding site-leucine-rich repeat genes reveal three anciently diverged classes with distinct evolutionary patterns. Plant Physiol., 170, 2095–2109.

Sievers, F., Wilm, A., Dineen, D., et al. (2011) Fast, scalable generation of high-quality protein multiple sequence alignments using Clustal Omega. Mol. Syst. Biol., 7, 539.

Sigrist, C.J.A., Castro, E. de, Cerutti, L., Cuche, B.A., Hulo, N., Bridge, A., Bougueleret, L. and Xenarios, I. (2013) New and continuing developments at PROSITE. Nucleic Acids Res., 41, D344–D347.

Smith, S.A. and Brown, J.W. (2018) Constructing a broadly inclusive seed plant phylogeny. Am. J. Bot., 105, 302–314.

Stamatakis, A. (2014) RAxML version 8: a tool for phylogenetic analysis and post-analysis of large phylogenies. Bioinformatics, 30, 1312–1313.

Stamatakis, A., Hoover, P. and Rougemont, J. (2008) A rapid bootstrap algorithm for the RAxML web servers. Syst. Biol., 57, 758–771.

Steuernagel, B., Jupe, F., Witek, K., Jones, J.D.G. and Wulff, B.B.H. (2015) NLR-parser: rapid annotation of plant NLR complements. Bioinformatics, 31, 1665–1667.

Steuernagel, B., Witek, K., Krattinger, S.G., et al. (2020) The NLR-annotator tool enables annotation of the intracellular immune receptor repertoire. Plant Physiol., 183, 468–482.

Tamborski, J. and Krasileva, K.V. (2020) Evolution of plant NLRs: from natural history to precise modifications. Annu. Rev. Plant Biol., 71, 355–378.

Toda, N., Rustenholz, C., Baud, A., Le Paslier, M.-C., Amselem, J., Merdinoglu, D. and Faivre-Rampant, P. (2020) NLGenomeSweeper: a tool for genome-wide NBS-LRR resistance gene identification. Genes, 11, 333.

Uehling, J., Deveau, A. and Paoletti, M. (2017) Do fungi have an innate immune response? An NLR-based comparison to plant and animal immune systems. PLOS Pathog., 13, e1006578.

Van Ghelder, C., Parent, G.J., Rigault, P., et al. (2019) The large repertoire of conifer NLR resistance genes includes drought responsive and highly diversified RNLs. Sci. Rep., 9, 11614.

Wang, Jizong, Hu, M., Wang, Jia, et al. (2019) Reconstitution and structure of a plant NLR resistosome conferring immunity. Science, 364, eaav5870.

Weber, L.M., Saelens, W., Cannoodt, R., Soneson, C., Hapfelmeier, A., Gardner, P.P., Boulesteix, A.-L., Saeys, Y. and Robinson, M.D. (2019) Essential guidelines for computational method benchmarking. Genome Biol., 20, 125.

Wickham, H., Averick, M., Bryan, J., et al. (2019) Welcome to the Tidyverse. J. Open Source Softw., 4, 1686.

Wu, C.-H., Abd-El-Haliem, A., Bozkurt, T.O., Belhaj, K., Terauchi, R., Vossen, J.H. and Kamoun, S. (2017) NLR network mediates immunity to diverse plant pathogens. Proc. Natl. Acad. Sci. U. S. A., 114, 8113–8118.

Wu, C.-H., Derevnina, L. and Kamoun, S. (2018) Receptor networks underpin plant immunity. Science, 360, 1300–1301.

Wu, C.-H., Krasileva, K.V., Banfield, M.J., Terauchi, R. and Kamoun, S. (2015) The “sensor domains” of plant NLR proteins: more than decoys? Plant-Microbe Interact., 6, 134.

Xie, J., Guo, G., Wang, Y., et al. (2020) A rare single nucleotide variant in *Pm5e* confers powdery mildew resistance in common wheat. New Phytol., 228, 1011–1026.

